# Early enhanced control of *Plasmodium yoelii* infection in IL-10–deficient mice is independent of IFN-γ, IL-12, and the humoral response

**DOI:** 10.64898/2026.03.23.713659

**Authors:** Meghan D. Jones, Kara A. O’Neal, Sheldon L. Zeltner, Alison R. Gouch, Jason S. Stumhofer

**Affiliations:** University of Arkansas for Medical Sciences, Department of Microbiology and Immunology, Little Rock, AR, United States of America

## Abstract

The outcome of a *Plasmodium* infection depends on the timely regulation of the robust pro-inflammatory response required to eliminate the parasite, but this response can cause tissue damage if not properly controlled. IL-10 is an important regulatory cytokine that prevents immunopathology during many *Plasmodium* infections; however, this protection comes at the expense of less effective parasite control. This is illustrated by infection with *P. yoelii*, in which mice exhibit a lower parasite load in the absence of IL-10. However, the immune components that limit parasite burden in the absence of IL-10 remain poorly understood. Abolishing IL-10 led to a predicted increase in T_H_1 polarization and higher production of IL-12 and IFN-γ. However, the enhanced production of these cytokines did not explain the improved parasite control seen in *Il10^-/-^* mice. Loss of IL-10 signaling reduced the accumulation of germinal center B cells and plasmablasts in the spleen, indicating a role for IL-10 in supporting the humoral response. However, although B cells are essential for survival, they do not play a critical role in early parasite control in IL-10–deficient mice. Moreover, *Il10^-/-^*mice lacking IFN-γ and B cells can limit early parasite expansion, suggesting that IL-10 suppresses host-protective pathways beyond the functions of B cells and IFN-γ in parasite control.

## Introduction

Malaria, caused by infection with *Plasmodium* parasites, remains a leading cause of death in many developing countries despite available therapeutics and the introduction of vaccines in some African countries (1, 2). Improving vaccine design by identifying new immune targets and enhancing protective immunity will be vital to reducing the global burden of malaria. Control of blood-stage *Plasmodium* infection is believed to involve two main strategies: a cell-mediated Type 1 helper (T_H_1) response and an antibody (Ab)–promoting T follicular helper (T_FH_) cell response, both of which are elicited in the spleen (3). In the T_H_1 response, T_H_1–polarized CD4^+^ T cells are the predominant source of IFN-γ during infection, although other cell types can also contribute to its production (4–6). IFN-γ, a pro-inflammatory cytokine, is linked to the recruitment, activation, and differentiation of inflammatory monocytes that promote parasite clearance through phagocytosis, production of free radicals, and other mediators in the phagolysosome (7–8). Alternatively, the T_FH_ response supports the activation, class switching, and differentiation of B cells into Ab-secreting cells through germinal center (GC) -independent and -dependent mechanisms. The Abs produced by this response are key for controlling and resolving blood-stage infections and for protecting against reinfection (9–11).

Juxtaposed to IFN-γ is the anti-inflammatory cytokine IL-10, which plays a crucial role in preventing immunopathology in numerous mouse models of autoimmune and infectious diseases, including malaria (6,12–15). IL-10 dampens inflammatory responses by downregulating pro-inflammatory cytokine production, co-stimulatory molecule expression, and major histocompatibility (MHC) molecule expression on antigen-presenting cells (APCs) (13, 15). Paradoxically, IL-10 can also promote hematological changes, including myelopoiesis, anemia, and thrombocytopenia when delivered to humans or under stress-inducing conditions, primarily by inducing T cell-mediated IFN-γ production (16–18). In the case of B cells, evidence supports a role for IL-10 in promoting survival, class switching, and Ab production (19–21). Conversely, IL-10 is also thought to limit GC reactions and humoral immunity in vivo (22–23). Regarding *Plasmodium* infection, evidence indicates that IL-10–producing B cells can limit pathology during infection (24, 25). However, the protection from immunopathology afforded by IL-10 comes at the expense of efficient parasite control. As a result, IL-10 production leads to increased parasite burdens during non-lethal and lethal *P. yoelii* infections (26, 27). This pattern extends to human infections with *P. vivax* and *P. falciparum*, where higher plasma IL-10 concentrations correlate with higher parasitemia (28–31). Therefore, understanding how IL-10, or the factors that regulate its expression, can be manipulated to enhance parasite control or vaccine-mediated immune responses without causing lethal or permanent tissue damage may influence future strategies for treating and preventing malaria.

Here, the impact of IL-10 on T and B cell responses during non-lethal *P. yoelii* 17X infection was investigated to determine correlates of protection arising in the absence of IL-10 signaling. Given the role of IL-10 in limiting pro-inflammatory cytokine production, it was hypothesized that a hyperbolic T_H_1 response with elevated IFN-γ production is necessary in IL-10–deficient mice to control the parasite more effectively. Findings here demonstrate that IL-10 limits control of infection but is not necessary to prevent severe tissue pathology. IL-10 modulates the host response by preventing prolonged T_H_1 polarization and is important in limiting splenic and systemic production of IL-12 and IFN-γ. However, these cytokines are not vital for protection in the IL-10–deficient mice. Furthermore, IL-10 promotes splenic GC and plasmablast responses but is not critical to elicit protective Ab responses. While Ab production is not vital for the enhanced parasite control in IL-10–deficient mice during early *P. yoelii* infection, B cells are ultimately critical for parasite clearance and survival. Moreover, even in the absence of B cells, IFN-γ production is not required for parasite control, suggesting that IL-10 suppresses alternative pathways that do not involve IFN-γ or B cells that can limit parasite expansion.

## Methods

### Mice and Infections

All animal studies were reviewed and approved by the University of Arkansas for Medical Sciences (UAMS) Institutional Animal Care and Use Committee (IACUC). Male and female wild-type (WT) C57BL/6J, *Il10^-/-^*, *μMT,* and *Ifngr1^-/-^* mice were purchased from The Jackson Laboratory, and *μMT*/*Il10^-/-^* and *Ifngr1^-/-^/Il10^-/-^* mice were generated in-house. All mice were housed and bred in specific-pathogen-free facilities at UAMS in accordance with institutional guidelines.

For non-lethal *Plasmodium yoelii* 17X (MR4, BEI Resources Repository) infection, parasitized red blood cells (pRBCs) from a cryopreserved stock were passaged intraperitoneally (i.p.) in a C57BL/6J mouse. Parasite burden was monitored using Giemsa-stained thin blood smears. Once parasitemia reached 1-5%, blood was collected, and 10^5^ pRBCs were injected i.p. to establish primary infection in experimental mice. All mice were infected between 8 and 12 weeks of age. Male and female mice were used to control for potential bias. For secondary infections, mice were challenged i.p. with 10^6^ *P. yoelii* 17X pRBCs 108 days p.i. Blood parasitemia during primary infections was determined by flow cytometry as previously described, beginning 5 days p.i. (32), while secondary infections were monitored using Giemsa-stained thin blood smears.

### Flow cytometry & surface staining

To generate single-cell suspensions for flow cytometry analysis, whole mouse spleens were processed through a 40 μm cell strainer (MIDSCI), and total splenocytes were subjected to RBC lysis (0.86% NH_4_Cl solution). Cells were kept in complete RPMI (RPMI 1640, 10% fetal bovine serum, 10% non-essential amino acids, 10% sodium pyruvate, 10% L-glutamine, 10% penicillin and streptomycin, and 1% 2-βME). Before staining, cells were washed in FACS buffer (1× PBS, 0.2% BSA, and 0.2% 0.5 M EDTA) and Fc receptors were blocked with anti-mouse CD16/32 (clone 2.4G2, BioXCell) in a buffer containing normal mouse and rat IgG (Invitrogen). For surface staining, all Abs were diluted in FACS buffer according to the manufacturer’s recommendations, and cells were incubated for 30 minutes at 4°C. Flow cytometry Abs used in the experiments are listed in Table S1. To minimize non-specific polymer interactions, any surface stain containing more than one Ab conjugated to a Brilliant Violet fluorophore was supplemented with 5 μL/sample of Super Bright Staining Buffer (Thermo Fisher Scientific). If no intracellular nuclear staining was required, cells were fixed in 1% paraformaldehyde (PFA; Electron Microscopy Sciences) for 10 minutes at room temperature. Following resuspension in FACS buffer, samples were acquired on a Cytek Northern Lights (Cytek Biosciences) spectral flow cytometer. All data were analyzed using FlowJo version 10 software. Fluorescence-minus-one (FMO) controls were used to set positive gates when necessary.

### Intracellular staining

For intracellular cytokine staining, processed splenocytes were restimulated by incubation with PMA (0.1 μg/mL) and ionomycin (1 μg/mL) in the presence of Brefeldin A (20 μg/mL) (Sigma-Aldrich) at 37°C for 4 hours. Following incubation, cells underwent surface staining, fixation in 1% PFA, and permeabilization in saponin (0.1%) diluted in FACS buffer. Abs against indicated cytokines (Table S1) were diluted in the permeabilization buffer, and cells were stained for 30 minutes at 4°C. Cells were then washed and resuspended in FACS buffer. Samples were acquired on a Cytek Northern Lights (Cytek Biosciences) spectral flow cytometer.

For T-bet and Bcl-6 staining, surface-stained splenocytes were fixed using the Foxp3/Transcription Factor Staining Buffer set (Invitrogen) per the manufacturer’s instructions. Briefly, cells were fixed in a buffer containing 1 part Fixation/Permeabilization Concentrate (Invitrogen) to 3 parts Fixation/Permeabilization Diluent (Invitrogen) for 30 minutes at 4°C. Abs were added at recommended dilutions in the Foxp3 Permeabilization/Wash buffer (1×; Invitrogen). Abs for flow cytometry are listed in Table S1. Cells were then washed twice and resuspended in FACS buffer. Samples were acquired on a Cytek Northern Lights (Cytek Biosciences) spectral flow cytometer.

### Hematological Analysis

To quantify total RBCs, hemoglobin, and hematocrit, 30 μL of blood was collected at the time of euthanasia in microtainer tubes containing K_2_EDTA (BD Biosciences). Samples were acquired on a Vetscan HM5 Hematology Analyzer (Zoetis). Blood from naïve mice from each genotype was used to assess the development of anemia during infection.

### H&E staining

Infected WT and *Il10^-/-^* mice were perfused through intracardiac injection of sterile PBS before removal of the liver and lungs. For livers, one lobe was extracted, placed in a tissue cassette (Epredia), and fixed in 10% formalin (Fisher Scientific). Lungs were inflated with 10% formalin through intratracheal injection before being placed in a cassette. Paraffin embedding, sectioning, and H&E staining were performed in collaboration with the Experimental Pathology Core at the University of Arkansas for Medical Sciences. Slides were imaged on an EVOS FL Auto 2 microscope (Life Technologies) at 10× magnification.

### Reticulocyte Imaging

Giemsa-stained thin blood smears were imaged on a Nikon Eclipse Ti2 (Nikon Instruments) confocal microscope using a Nikon Digital Sight DS-Fi2 (Nikon Instruments) camera. Images were taken at 100× magnification under oil immersion.

### Antibody ELISAs

For Ab ELISAs, 96-well F-bottom microplates (Greiner) were coated overnight at 4°C with recombinant *P. yoelii* MSP-1_19_ protein (2.5 μg/mL) diluted in sodium carbonate buffer (8.4% sodium bicarbonate, 10.4% sodium carbonate in diH_2_O). Plates were washed (0.05% Tween in 1× PBS) and blocked with 5% Fetal plex (Gemini Biosciences) in PBS for 1 hour at 37°C. Serum was diluted 1:50 or 1:100 for IgM detection and 1:50 or 1:500 for IgG detection, then serially diluted 1:3 in 5% Fetal plex in PBS. Plates were incubated at 37°C for 2 hours. HRP-conjugated IgM (1:4000) or IgG (1:6000) (Southern Biotech) diluted in 5% Fetal plex in PBS was added, and plates were incubated for 1 hour at 37°C. KPL SureBlue substrate (SeraCare) was used for detection. The enzyme reaction was allowed to proceed for 5 minutes before the KPL TMP stop solution (SeraCare) was added. Plates were read at an absorbance of 450 nm on a CLARIOStar plate reader (BMG Labtech).

### Cytokine ELISAs and multiplex bead assay

A bead-based LEGENDplex™ assay (BioLegend, Cat No. 740150) was used according to the manufacturer’s instructions to determine the serum concentrations of selected cytokines, including IFN-γ, TNF-α, and IL-6. Serum was diluted 1:2 in the kit’s assay buffer. Samples were transferred to FACS tubes and acquired on an LSRII Fortessa (Becton Dickson) flow cytometer. LEGENDplex data analysis software (BioLegend) was used to generate standard curves and estimate cytokine concentrations.

To determine the concentration of serum IL-12p40, 96-well F-bottom microplates (Greiner) were coated overnight with anti-IL-12p40 (clone C15.6, BioLegend) diluted in PBS at 4 °C. Plates were washed (0.05% Tween in 1× PBS) and blocked for 1 hour at 37°C with 5% Fetal Bovine Serum (FBS) (R&D Systems) in PBS. Protein standards were diluted two-fold in cRPMI, starting at 20 ng/mL. Protein standards and serum samples (40 μL/well) were incubated for 2 hours at 37°C. Secondary biotinylated anti-IL-12p40 (clone C17.8, BioLegend) was added to plates diluted (1:1000) in ELISA buffer (0.1% BSA, 0.5% Tween 20 in 1× PBS). Plates were incubated at room temperature for 1 hour. HRP-conjugated streptavidin was diluted (1:2000) in ELISA buffer before addition to the plate. Plates were incubated at room temperature for 30 minutes. ABTS (Life Technologies) was used for detection, and plates were developed for 10 minutes. Plates were read at an absorbance of 405 nm on a CLARIOStar plate reader (BMG Labtech). ClarioStar MARS software v3.40 R2 (BMG Labtech) was used to generate a standard curve from absorbance values obtained from serially diluted protein standards. This standard curve was used to estimate IL-12p40 concentrations in the serum samples.

### ELISpots

Mixed cellulose ester filter plates (Millipore) were activated with 35% EtOH for 1 minute and coated with recombinant MSP-1_19_ (2.5 µg/mL) diluted in sterile PBS overnight at 4°C. The following day, plates were blocked for 30 minutes at room temperature with 5% FBS (R&D Systems) in PBS. Splenocytes were added (200,000/well) in cRPMI made with FBS. Plates were incubated at 37°C in 5% CO_2_ for 24 hours. Plates were then blocked with 5% FBS (R&D Systems) diluted in PBS for 1 hour at 37°C. Secondary goat anti-mouse IgM-AP (Southern Biotech) (1:1000) or goat anti-mouse IgG-AP (Southern Biotech) (1:800) diluted in 5% FBS (R&D Systems) in PBS was added for 1 hour at room temperature. Plates were developed for 30 minutes with 5-bromo-4-chloro-3-indolyl-phosphate (BCIP) (0.175 mg/mL) and nitroblue tetrazolium (NBT) (0.225 mg/mL) diluted in a developing buffer (2.1 g NaHCO_3_, 50.8 mg MgCl_2_, and 250 mL H_2_0). The plates were rinsed with water and allowed to dry for 24 hours. Plates were read on an AID ELISpot reader, and spots were enumerated using the AID ELISpot software version 7.0 (AID GmbH).

### *In vivo* antibody blockade

To determine the importance of IFN-γ signaling without IL-10 during infection, female WT mice were i.p. administered either 200 μg of Rat IgG (Sigma-Aldrich), 500 μg of anti-IFN-γ (clone XMG1.2, BioXcell), 200 μg of anti-IL-10R (clone 1B1.3A, BioXcell), or both anti-IFN-γ and anti-IL-10R diluted in sterile PBS on day 1, 4, 7, 10, 13, and 16 p.i. To determine the necessity of IL-12 and/or IFN-γ without IL-10 during infection, *Il10^-/-^* mice were i.p. administered either 500 μg of Rat IgG (Sigma-Aldrich), 500 μg of anti-IL-12p75 (clone R2-9A5, BioXcell), or 500 μg of anti-IFN-γ (clone XMG1.2, BioXcell) diluted in sterile PBS on day 0, 3, 6, 9, and 12 p.i. To determine the importance of IFN-γ when both IL-10 and B cells are absent during infection, male and female *μMT*/*Il10^-/-^* mice were i.p. administered either 500 μg of Rat IgG (Sigma-Aldrich) or 500 μg of anti-IFN-γ (clone XMG1.2, BioXcell) diluted in sterile PBS on day 0, 3, 6, 9, and 12 p.i.

### Statistical analysis

GraphPad Prism 10 (GraphPad Software, Inc., San Diego, CA) was used for statistical analysis. Statistical significance was determined using a nonparametric Mann-Whitney U test for single-group comparisons. For multiple comparisons between groups, statistical significance was determined using a Kruskal-Wallis test with Dunn’s post hoc test. For multiple comparisons between two variables, statistical significance was assessed using two-way ANOVA with a Holm-Sîdak post-hoc test. Comparisons of survival curves were performed using a Mantel-Cox test. Statistical comparisons were made to the WT control group. A *p*-value of < 0.05 determined significance. Additional details on specific statistical tests are provided in the figure legends.

## Results

### Eliminating global IL-10 production improves control of *P. yoelii* infection without eliciting lethal immunopathology

Previous observations suggest that IL-10 limits parasite control during *P. yoelii* 17X infection when mice are subjected to 10^4^ pRBCs intravenously (i.v.) (26). Raising the parasite inoculum and changing the route of infection did not alter this phenotype (**Fig. 1A, B**). During *P. yoelii* infection, *Il10^-/-^* mice have significantly lower parasite burdens compared to WT mice; however, peak parasitemia (day 14-16 p.i) and the time of clearance (day 24-26 p.i.) are invariant in the absence of IL-10 (**Fig. 1A, B**). Given the preference of *P. yoelii* for invading reticulocytes, (33) *Il10^-/-^* mice were examined for a defect in reticulocyte production before and after infection. Notably, the lower parasite burdens in *Il10^-/-^* mice are not due to inadequate reticulocyte availability before infection (**Fig. S1A-C**). However, a drop in reticulocyte availability was observed at day 7 in *Il10*^-/-^ mice (**Fig. 1C**), leading to a shift in RBC invasion preference toward normocytes (**Fig. S1C**). This dip in reticulocytes was temporary, as their abundance thereafter increased at a rate similar to that of WT mice, and parasite preference shifted back to reticulocytes as their availability increased. While reticulocyte abundance increased from day 9 onward, correlating with a steady rise in parasitemia in WT mice, parasitemia did not rise to the same extent in the *Il10^-/-^*mice (**Fig. S1 A-B**). Thus, a defect in reticulocytosis does not account for the lack of parasite expansion in *Il10*^-/-^ mice.

**Fig 1.**
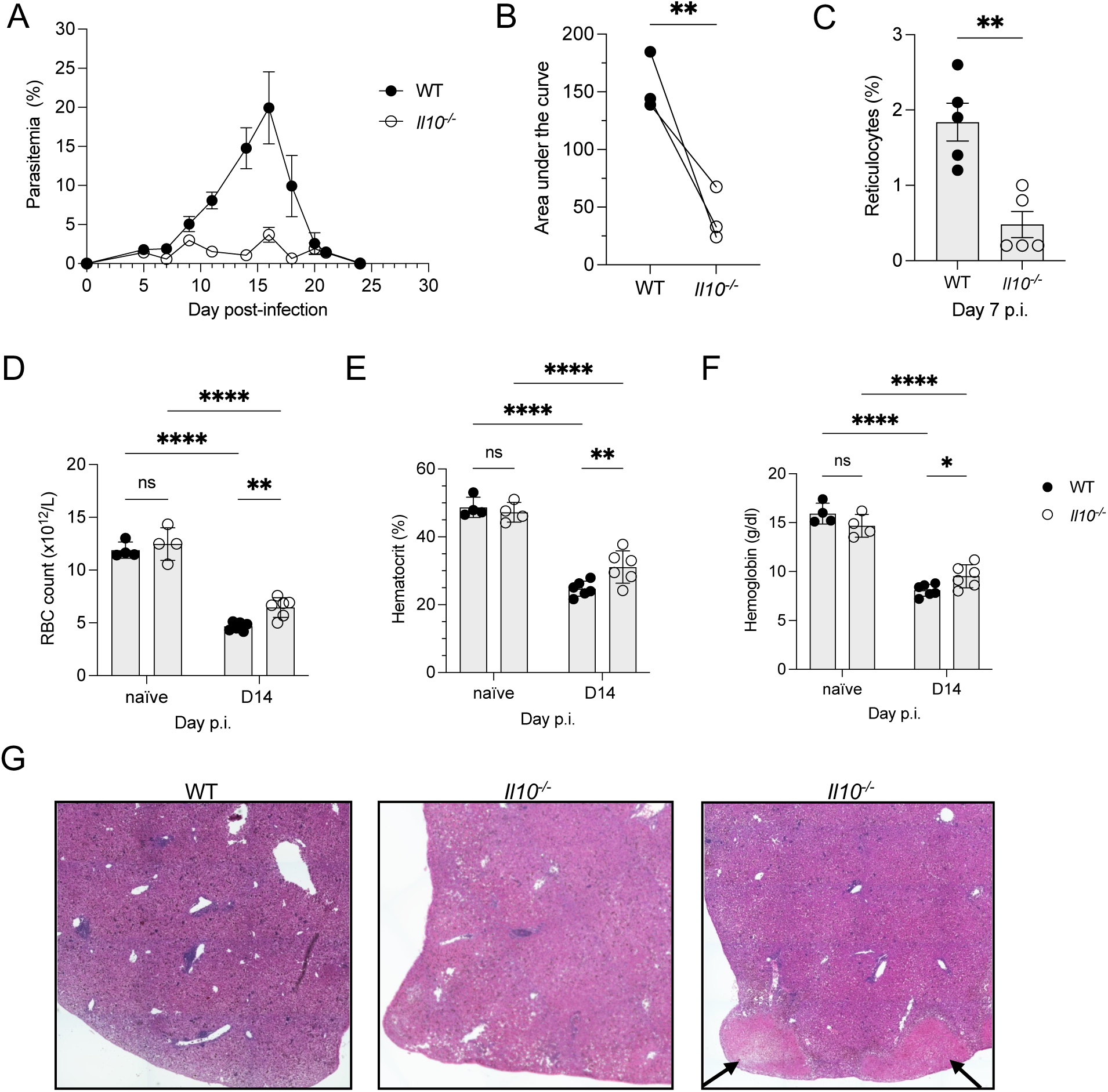
IL-10 limits control of *P. yoelii* infection without eliciting severe immunopathology. WT and *Il10^-/-^* mice were infected i.p. with 10^5^ *P. yoelii* 17X pRBCs. (**A**) Representative parasitemia curve during primary infection. Parasitemia was monitored by flow cytometry starting on day 5 p.i. The data represent 3 independent experiments, with 4-6 mice per group. (**B**) The area under the curve was measured for the entire curve for WT and *Il10^-/-^*mice from 3 independent experiments. (**C**) Comparison of reticulocyte frequency at day 7 p.i. determined through Giemsa-stained thin blood smears. (**D**) RBC count, (**E**) hemoglobin, and (**F**) hematocrit values were determined by analysis of blood collected from naïve mice and on D14 p.i. using a Vetscan Hematology Analyzer. (**G**) H&E-stained liver sections from D14 p.i. Microscopic images were taken at 10× magnification. Arrows denote areas of necrosis and fibrosis. (**B-C**) A non-parametric Mann-Whitney t-test and (**D-F**) a two-way ANOVA determined significance with a post hoc Holm-Sidak’s multiple comparisons test. \**p* < 0.05, ** *p* < 0.01, **** *p* < 0.0001, ns = non-significant.

Anemia is a common, albeit potentially lethal, consequence of *Plasmodium* infection (34, 35) and triggers the production of new RBCs, as evidenced by an increase in reticulocytes. To determine the severity of anemia during infection in the absence of IL-10, blood was collected at days 7 and 14 p.i., and total RBCs, hemoglobin, and hematocrit were quantified. Signs of anemia were present in all mice at day 7 p.i., as indicated by lower RBC counts, hematocrit, and hemoglobin values compared with naïve mice of the respective genotype (**Fig. S1D-F**), though no differences were noted between groups. However, by day 14 p.i., signs of anemia were more pronounced in WT mice, even though both groups were considered anemic (**Fig. 1D-F**).

In several mouse models, IL-10 deficiency exacerbates tissue pathology through an enhanced inflammatory response, leading to decreased survival (6, 36–37). However, in this case, *Il10^-/-^* mice survive infection with *P. yoelii*, though tissue pathology could still occur. Thus, it was of interest to determine if immunopathology was present in the *Il10^-/-^*mice during infection. The liver is a common site of pathology during *Plasmodium* infection (34, 38). Therefore, livers were harvested, and sections were stained with H&E to detect pathological changes. At day 14 p.i., WT mice show no signs of severe liver pathology (**Fig. 1G**). In the absence of IL-10, tissue pathology was inconsistent, with evidence of tissue necrosis in a few, but not all, liver sections (**Fig. 1G**). Furthermore, lung sections were collected at day 14 p.i. and subjected to H&E staining. In this case, there was no evidence of enhanced immunopathology in the IL-10–deficient lungs (**Fig. S1G**). Thus, this data, consistent with other (26), suggests that abrogating IL-10 production during *P. yoelii* infection improves parasite control without inducing lethal tissue pathology or severe anemia. How infection is more efficiently controlled in the absence of IL-10 signaling remains unknown.

### The splenic T_H_1 response is elevated in the absence of IL-10 during *P. yoelii* infection

To define the correlates of protection that arise in the absence of IL-10 signaling, changes in the T_H_1 response were investigated. IL-10 is known to dampen the T_H_1 response indirectly by restricting proinflammatory cytokine production by innate cells and by halting antigen presentation by downregulating MHC expression on APCs (15). Therefore, it was hypothesized that eliminating IL-10 signaling would enhance the T_H_1 response during infection, thereby improving parasite control. To determine changes in the T_H_1 response in the absence of IL-10, spleens from WT and *Il10^-/-^* mice were harvested at day 5 p.i. to quantify CD4^+^ T cell populations. The total number of splenocytes and CD4^+^ T cells was similar between groups (**Fig. S2A,B**). The frequency of antigen-experienced CD4^+^ T cells (CD11a^hi^CD44^hi^) was significantly lower in *Il10^-/^*^-^ mice; however, their total numbers were similar (**Fig. S2C-E**). To measure changes in T_H_1 polarization, T-bet and CXCR6 expression were quantified. Expression of this chemokine receptor is associated with T_H_1 cells (39) and, more recently, implicated as a marker of regulatory Type 1 (Tr1) cells (40). At day 5 p.i., T-bet and CXCR6 expression on antigen-experienced CD4^+^ cells was not impacted by IL-10 (**Fig. 2A-F**). Thus, this suggests that IL-10 does not constrain the initial polarization of T_H_1 cells during infection.

**Fig 2.**
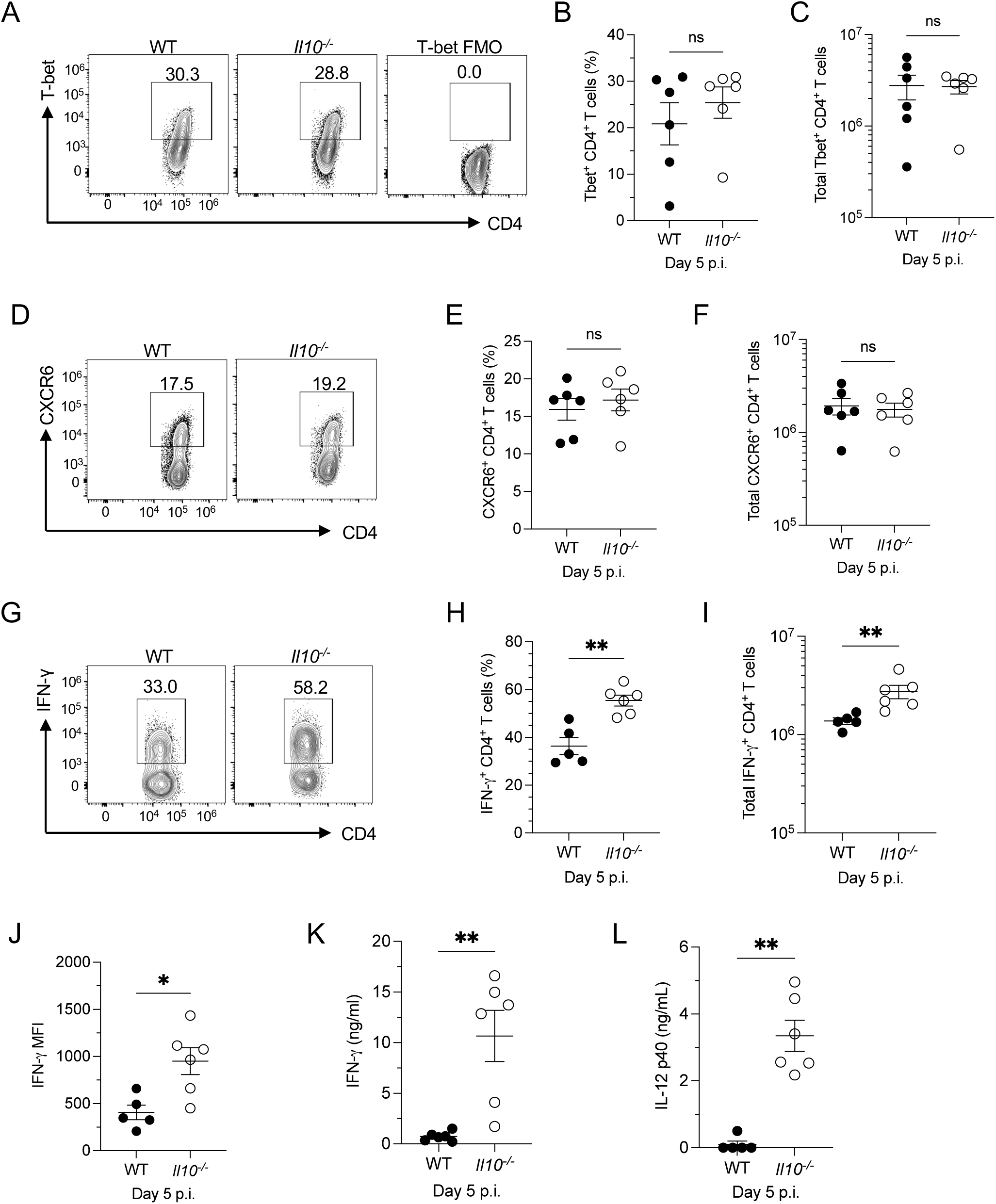
IL-10 suppresses IFN-γ production early in infection but does not affect the initiation of T_H_1 polarization during *P. yoelii* infection. WT and *Il10^-/-^* mice were infected i.p. with 10^5^ *P. yoelii* 17X pRBCs. (**A**) Representative flow plots of live, single-cell, T-bet^+^ CD4^+^ T cells in the spleen at day 5 p.i. FMO (Fluorescence minus one) controls were used to set gates. (**B**) Frequency of T-bet^+^ CD4^+^ T cells out of total CD11a^hi^CD44^hi^ CD4^+^ T cells. (**C**) Total T-bet^+^ CD11a^hi^CD44^hi^ CD4^+^ T cells. (**D**) Representative flow plots of live, single-cell, CXCR6^+^ CD4^+^ T cells in the spleen at day 5 p.i. (**E**) Frequency of CXCR6^+^ out of CD11a^hi^CD44^hi^ CD4^+^ T cells. (**F**) Total CXCR6^+^ CD11a^hi^CD44^hi^ CD4^+^ T cells in the spleen. (**G**) Representative flow plots of live, single-cell, IFN-γ^+^ CD4^+^ T cells at day 5 p.i. in the spleen. (**H**) Frequency of IFN-γ^+^ CD4^+^ T cells out of total CD4^+^ T cells. (**I**) Total IFN-γ^+^ CD4^+^ T cells in the spleen. (**J**) Median fluorescent intensity (MFI) of IFN-γ in CD4^+^ T cells. (**K**) Serum concentration of IFN-γ at day 5 p.i. determined by Biolegend^TM^ multiplex assay. (**L**) Serum concentration of IL-12p40 at day 5 p.i. determined by ELISA. (**A-J**) The data represent 2 independent experiments with 4-6 mice per group. (**B, C, E, F, H, I, J, K, L**) A non-parametric Mann-Whitney t-test determined significance. \**p* < 0.05, ** *p* < 0.01, ns = non-significant.

IFN-γ, produced by T_H_1–polarized CD4^+^ T cells, plays an integral role in controlling *Plasmodium* infections (7,8). Though there was no elevated CXCR6 or T-bet expression at day 5 p.i., there was significantly more IFN-γ^+^ CD4^+^ T cells in the *Il10^-/-^* mice (**Fig 2G-I**). The median-fluorescent intensity (MFI) of IFN-γ in CD4^+^ T cells was significantly elevated in the *Il10^-/-^* mice at day 5 p.i., as well, suggesting more IFN-γ production on a per-cell basis (**Fig 2J**). IFN-γ production was also elevated in the *Il10^-/-^*CD8^+^ T cells at this time; the MFI was comparable between groups (**Fig S2F-I**). Along with the elevation of IFN-γ in the spleen, systemic production of IFN-γ was enhanced in the *Il10^-/-^* mice (**Fig 2K**). To further characterize changes in the T_H_1 response, serum IL-12, a cytokine known to induce T_H_1 polarization (41), was measured. As expected, systemic production of IL-12p40 was elevated in the *Il10^-/-^* mice at day 5 p.i. (**Fig 2L**). Other pro-inflammatory cytokines, such as TNF-α and IL-6, were also measured in serum on day 5 p.i. While there was no significant change in IL-6 production in the absence of IL-10, TNF-α was significantly elevated in the *Il10^-/-^* mice (**Fig. S2J,K**). Thus, these data suggest that early in infection, IL-10 suppresses IFN-γ, IL-12, and TNF-α production.

To characterize how the absence of IL-10 impacts the sustainability of the T_H_1 response during *P. yoelii* infection, the T_H_1 response was evaluated at day 14 p.i., a time that corresponds with or just before peak parasitemia in WT and *Il10^-/-^*mice (**Fig. 1A**). Similar to day 5, total splenocytes and CD4^+^ T cells were comparable between groups (**Fig. S3A,B**). Although their frequencies were similar, fewer antigen-experienced CD4^+^ T cells were present in *Il10^-/^*^-^ mice at day 14 (**Fig. S3C-E**). However, T-bet and CXCR6 expression were significantly elevated amongst the antigen-experienced *Il10^-/^*^-^ CD4^+^ T cells (**Fig. 3A-F**). Thus, these results suggest that IL-10 suppresses CXCR6 and T-bet expression, and that in its absence, T_H_1 polarization is sustained.

**Fig 3.**
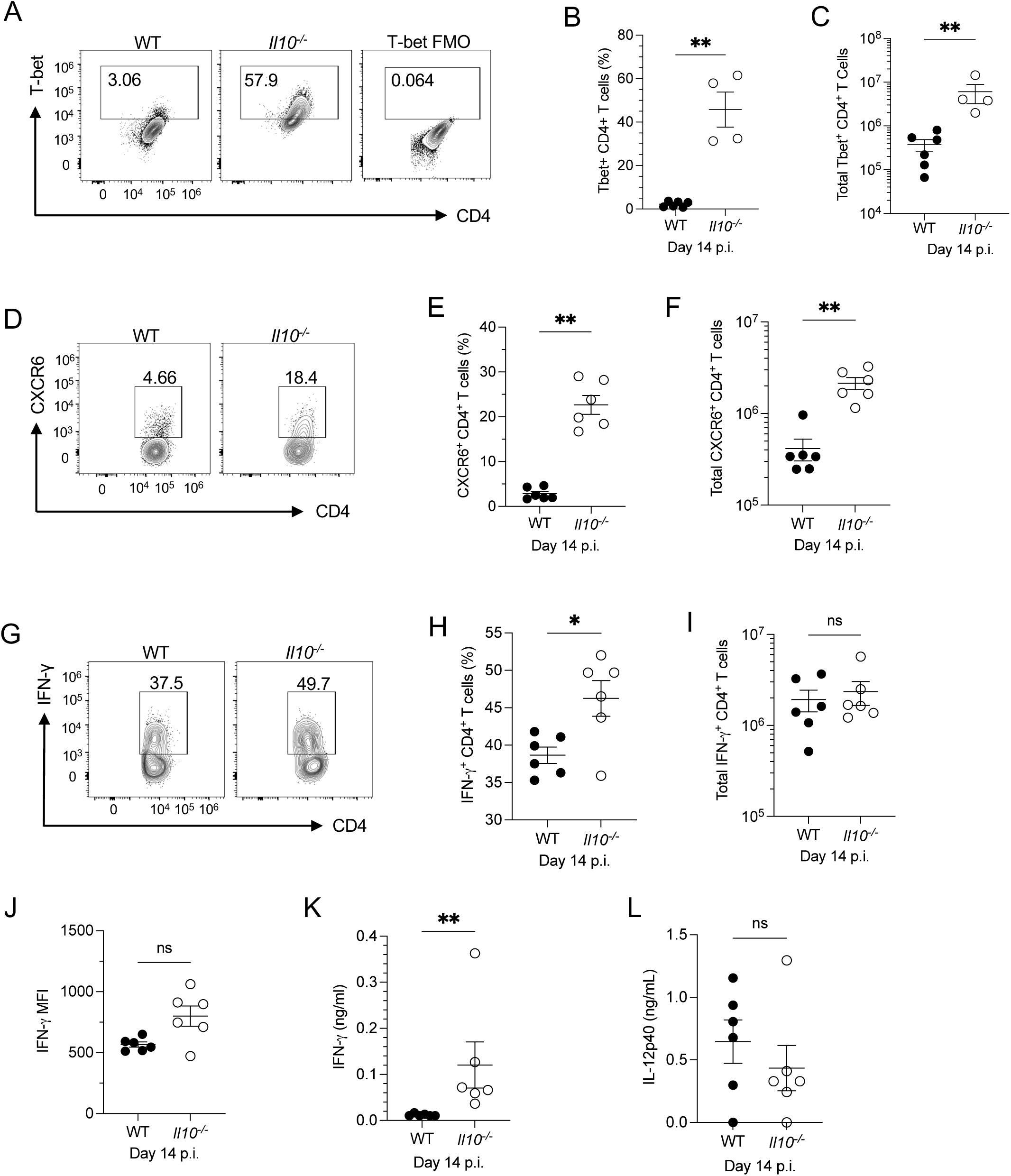
IL-10 suppresses T_H_1 polarization during *P. yoelii* infection. WT and *Il10^-/-^* mice were infected i.p. with 10^5^ *P. yoelii* 17X pRBCs. (**A**) Representative flow plots of live, single-cell, T-bet^+^ CD4^+^ T cells in the spleen at day 14 p.i. FMO controls were used to set gates. (**B**) Frequency of T-bet^+^ CD4^+^ T cells out of total CD11a^hi^CD44^hi^ CD4^+^ T cells. (**C**) Total T-bet^+^ CD11a^hi^CD44^hi^ CD4^+^ T cells. (**D**) Representative flow plots of live, single-cell, CXCR6^+^ CD4^+^ T cells in the spleen at day 14 p.i. (**E**) Frequency of CXCR6^+^ out of CD11a^hi^CD44^hi^ CD4^+^ T cells. (**F**) Total CXCR6^+^ CD11a^hi^CD44^hi^ CD4^+^ T cells in the spleen. (**G**) Representative flow plots of IFN-γ^+^ CD4^+^ T cells at day 14 p.i. in the spleen. (**H**) Frequency of IFN-γ^+^ CD4^+^ T cells out of total CD4^+^ T cells. (**I**) Total IFN-γ^+^ CD4^+^ T cells in the spleen. (**J**) Median fluorescent intensity (MFI) of IFN-γ in CD4^+^ T cells. (**K**) Serum concentration of IFN-γ at day 14 pi. determined by Biolegend^TM^ multiplex assay. (**L**) Serum concentration of IL-12p40 at day 14 p.i. determined by ELISA. (**A-J**) The data are representative of 2 independent experiments with 4-6 mice per group. (**B, C, E, F, H, I, J, K, L**) A non-parametric Mann-Whitney t-test determined significance. \**p* < 0.05, ** *p* < 0.01, ns = non-significant.

The heightened number of T_H_1–polarized CD4^+^ T cells correlated with a significant increase in the frequency of IFN-γ^+^ CD4^+^ T cells in the spleen of *Il10^-/^*^-^ mice, but total numbers were comparable between groups (**Fig. 3G-I**). Similarly, IFN-γ–producing CD8^+^ T cells were comparable between groups on day 14 p.i. (**Fig. S3F-H**). While the MFI of the IFN-γ^+^ CD4^+^ T cells was elevated in *Il10^-/^*^-^ mice, the difference was not significant compared with WT mice (**Fig. 3J**). Alternatively, the MFI of IFN-γ^+^ CD8^+^ T cells from *Il10^-/^*^-^ mice was significantly reduced at this time compared with WT mice (**Fig. S3I**). Furthermore, the concentration of IFN-γ remained significantly elevated in the serum of *Il10^-/^*^-^ mice through day 14 p.i. However, the concentration of IFN-γ was lower at day 14 than at day 5 in both sets of mice (**Fig 3K**; **Fig 2K**).

The concentration of IL-12 was elevated in the serum of *Il10^-/^*^-^ mice at day 10 (**Fig. S3J**) but not day 14 p.i (**Fig. 3L**). Similarly, by day 14 p.i., IL-10 had no significant impact on TNF-α production, and IL-6 production was negligible in both genotypes (**Fig S3K,L**). These results suggest that IL-12, TNF-α, and IFN-γ production can be restrained by additional mechanisms in the absence of IL-10, which could explain why *Il10^-/^*^-^ mice do not succumb to severe immunopathology in response to *P. yoelii* infection. Although IL-10 signaling appears to be critical for reducing the overall T_H_1 response, which may contribute to the enhanced parasite control in *Il10^-/^*^-^ mice.

### IL-10 supports the plasmablast and germinal center responses during *P. yoelii* infection

A strong humoral response, including *Plasmodium*-specific Ab production, is needed to control *P. yoelii* infection (9–11). Therefore, it was of interest to determine whether the humoral response is enhanced in the absence of IL-10, potentially contributing to improved parasite control. To allow time for the humoral response to develop, spleens were harvested at days 7 and 14 p.i. to quantify cell populations. Total splenocytes at day 7 p.i. were comparable between WT and *Il10^-/-^* mice (**Fig. S4A**), but plasmablasts were significantly reduced in the absence of IL-10 at day 7 p.i. (**Fig. 4A-C**). However, by day 14 p.i., plasmablast frequency and numbers were similar in all mice (**Fig. S4B-D**), indicating that any requirement for IL-10 in promoting plasmablast accumulation is transient.

**Fig 4.**
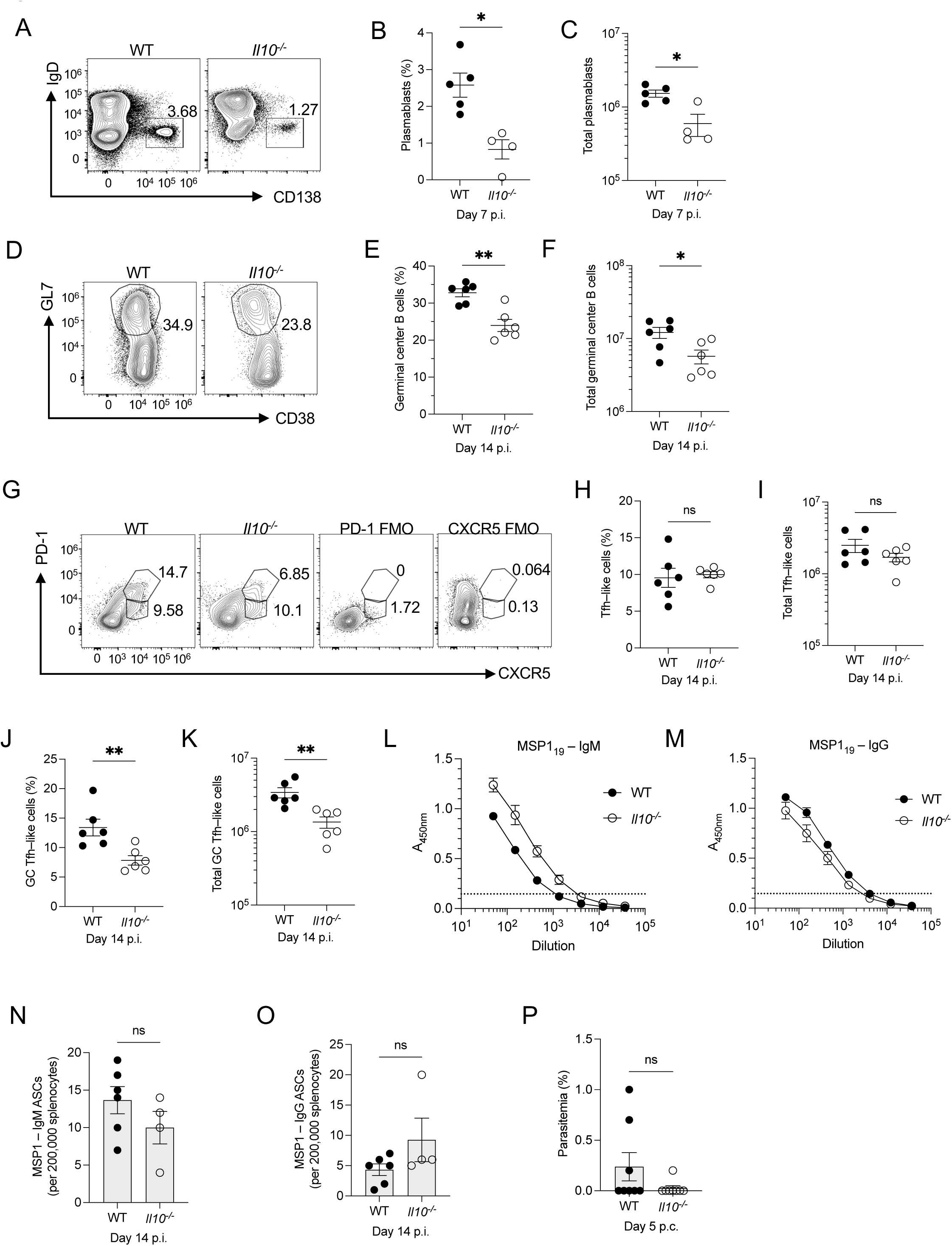
IL-10 promotes plasmablast and germinal center responses but is not required to elicit protective immunity during *P. yoelii* infection. WT and *Il10^-/-^*mice were infected i.p. with 10^5^ *P. yoelii* 17X pRBCs. (**A**) Representative flow plots of live, single-cell, CD138^+^IgD^-^splenic plasmablasts 7 days p.i. (**B**) Frequency of plasmablasts out of CD3^-^, Ter119^-^, CD11c^-^, CD11b^-^ cells. (**C**) Total plasmablasts in the spleen. (**D**) Representative flow plots of splenic live, single-cell, CD38^-^GL7^+^ GC B cells at 14 days p.i. (**E**) Frequency of splenic GC B cells out of total CD19^+^B220^+^ B cells. (**F**) Total GC B cells in the spleen. (**G**) Representative flow plots of live, single-cell, PD-1^Int^CXCR5^+^ Tfh-like cells and PD-1^hi^CXCR5^+^ GC Tfh cells in the spleen at day 14 p.i. FMO controls were used to set gates. (**H**) Frequency of Tfh-like cells out of CD11a^hi^CD44^hi^ CD4^+^ T cells. (**I**) Total Tfh-like cells in the spleen. (**J**) Frequency of GC Tfh cells out of CD11a^hi^CD44^hi^ CD4^+^ T cells. (**K**) Total GC Tfh cells in the spleen. (**L**) Serum IgM and (**M**) IgG titers against MSP-1 at day 14 p.i. detected by ELISA. (**N**) Total ASCs producing MSP-1– IgM and (**O**) MSP-1– IgG in the spleen at day 14 p.i. determined by ELISpot. (**P**) WT and *Il10^-/-^* mice were challenged 108 days after the primary infection with 10^6^ *P. yoelii* 17X pRBCs. Parasitemia was determined 5 days post-challenge through Giemsa-stained thin blood smears. (**A-P**) The data are representative of 2 independent experiments with 4-6 mice per group. (**B, C, E, F, H-K, M, O, P**) A non-parametric Mann-Whitney t-test determined significance. \**p* < 0.05, ** *p* < 0.01, ns = non-significant. The dotted line in **L** and **M** represents the limit of detection.

Given the importance of the GC response in protection against *Plasmodium* infection (9–11) and the role IL-10 has in supporting this response (42–43), GC B cells and T_FH_ cells were examined. GC B cells were detectable in both groups of mice by day 7 p.i. (**Fig. S4E-G**). Though the percentage of GC B cells was reduced in *Il10^-/-^* mice, this did not impact their overall numbers. However, while this population expanded by day 14 p.i., the frequency and total number of CD38^-^GL7^+^ GC B cells were significantly lower in *Il10^-/-^*mice (**Fig. 4D-F**). Examination of T_FH_ cell populations indicated that T_FH_-like cells (PD-1^+^CXCR5^+^) were not impacted by the loss of IL-10 at day 7 and 14 p.i. (**Fig. 4G-I; Fig. S4H-J**). In contrast, GC T_FH_ cell populations, defined by high PD-1 and CXCR5 expression, were comparable on day 7 p.i. (**Fig S4H, K-L**), but were significantly reduced in the *Il10^-/-^* mice at day 14 p.i. (**Fig 4G, J-K**). These results suggest that the GC response is constrained in the absence of IL-10, supporting previous observations of decreased splenic GC B cells when B cells lack IL-10R during *P. yoelii* infection (42).

Given the defects in the cell populations associated with the humoral response, it was of interest to assess whether the loss of IL-10 affected *Plasmodium*-specific Ab production and protection after secondary infection. Serum Ab titers against the *Plasmodium* antigen MSP-1 were measured using ELISAs. At day 14 p.i., the *Il10^-/-^* mice had similar titers of MSP-1–specific IgM and IgG compared to WT mice (**Fig. 4L, M**). Also, the accumulation of antibody-secreting cells (ASCs) producing MSP-1–specific IgM and IgG in the spleen was not impacted by the loss of IL-10 at day 14 p.i. (**Fig. 4N, O**). Following clearance of the infection, IgM and IgG titers specific to MSP-1, as well as MSP-1–specific ASCs, remained similar across all mice (**Fig. S4M, O**). Thus, the loss of IL-10 did not significantly impact parasite-specific Ab production. To determine whether Abs and the memory response generated during primary infection in the absence of IL-10 are functional and protective, WT and *Il10^-/-^* mice were challenged with a higher inoculum of *P. yoelii* 17X pRBCs, and parasitemia was monitored. The majority of the *Il10^-/-^* mice had no detectable parasites by day 5 post-challenge (p.c.) (**Fig 4P**), and all mice, WT and *Il10^-/-^*, were clear of parasites by day 7 p.c. (*data not shown*). Taken together, the data suggest that IL-10 supports, rather than limits, the plasmablast and GC responses. Furthermore, the constrained humoral response in *Il10^-/-^*mice did not have any short or long-term effects on host immunity.

### B cells are essential for survival but not early parasite control in *Il10^-/-^* mice during *P. yoelii* infection

Genetically mutated mice lacking mature B cells (*μMT*) cannot control *P. yoelii* infection due to defects in the humoral response (9, 44), emphasizing the importance of B cells in controlling this infection. To further elucidate whether the humoral response contributes to the improved parasite control observed in *Il10^-/-^* mice, *μMT*/*Il10^-/-^* mice were generated. Early in infection, parasitemia was comparable between all groups of mice (**Fig. 5A**). By day 9 p.i., the *Il10^-/-^* and *μMT*/*Il10^-/-^* mice had significantly lower parasite burdens compared to the WT and *μMT* controls (**Fig. 5A, B**), with parasite burdens in the *μMT* mice closer to that of the WT mice. The *Il10^-/-^* and *μMT*/*Il10^-/-^* mice had comparable parasitemia through day 15 p.i. However, after this point, the *μMT*/*Il10^-/-^* mice began to lose control of the infection, whereas *Il10^-/-^* mice cleared it by day 21 (**Fig 5A**). Similarly, the parasitemia of the *μMT* mice eclipsed that of the WT mice on day 15, suggesting that B cells and Abs are needed at this point to restrain the parasite burden, regardless of the presence of IL-10. Compared with the *μMT* mice, the *μMT*/*Il10^-/-^*mice maintained better control of parasitemia throughout infection, but they still succumbed to infection (**Fig 5C**). Thus, these results reinforce the need for B cells to control and clear *P. yoelii* infection, particularly after two weeks post-infection. Although the Ab response is not impacted by the loss of IL-10, it ultimately contributes to the ability of *Il10*^-/-^ mice to control and clear the infection. Also, the results suggest that additional aspects of the immune response, beyond the humoral response, are responsible for controlling early parasite burden in the absence of IL-10.

**Fig 5.**
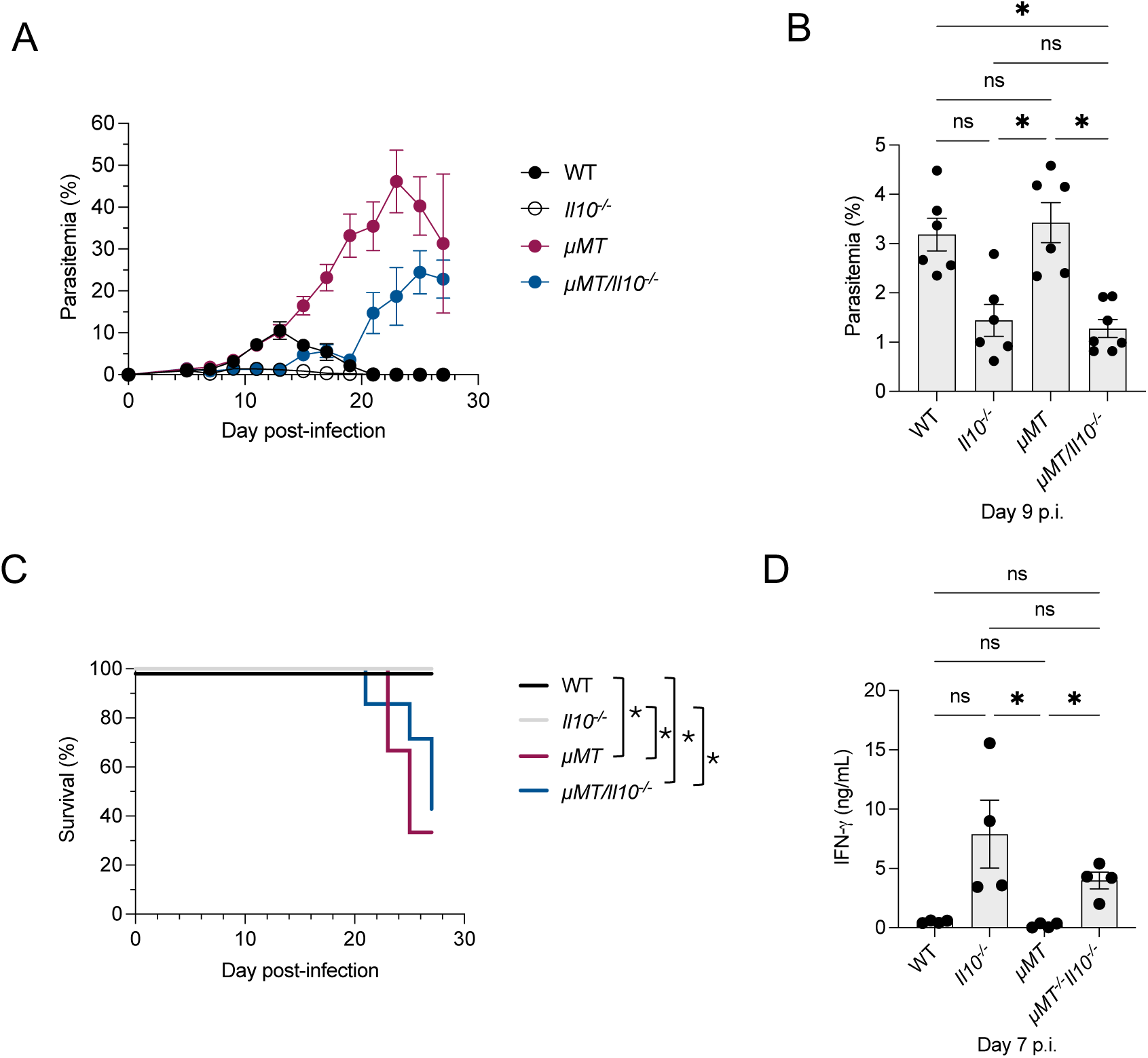
B cells are required for survival of *P. yoelii* infection but are not required for early parasite control in the absence of IL-10. WT, *Il10^-/-^*, *μMT*, and *μMT*/*Il10^-/-^* mice were infected i.p. with 10^5^ *P. yoelii* 17X pRBCs. (**A**) Representative parasitemia curve during primary infection. Parasitemia was monitored by flow cytometry starting at day 5 p.i. The data are representative of 2 independent experiments, with 3-5 mice per group. (**B**) Comparison of parasitemia measured at day 9 p.i. (**C**) Representative survival curve for the primary infection. Mice were euthanized when the parasite burden exceeded 50% or when prominent signs of morbidity appeared. (**D**) Serum concentration of IFN-γ at day 7 p.i. determined by ELISA. (**B, D**) A non-parametric Kruskal-Wallis test with a post hoc Dunn’s multiple comparisons test and (**C**) a Mantel-Cox test determined significance. \**p* < 0.05, ** *p* < 0.01, ns = non-significant.

### Enhanced IFN-γ and IL-12 production is not required to improve parasite control in the absence of IL-10 signaling

The sustained, enhanced parasite control in the *μMT*/*Il10^-/-^* mice compared to the WT and *μMT* mice over the first two weeks of infection suggests that factors other than B cells contribute to parasite control in the absence of IL-10 (**Fig. 5A,B**). One such candidate is IFN-γ, as it is also elevated in the serum at day 7 in the *μMT*/*Il10^-/-^* mice (**Fig. 5D**). To determine if increased IFN-γ production in *Il10^-/-^* mice was required to maintain this protection, IL-10R and IFN-γ were blocked with monoclonal Abs in WT mice during *P. yoelii* infection. Similar to the *Il10^-/-^* mice, blocking IL-10R resulted in significantly lower parasitemia during infection compared to the Rat IgG control (**Fig. 6A,B**). Co-administration of anti-IFN-γ and anti-IL-10R did not alleviate parasite control compared to the anti-IL-10R group. (**Fig. 6A,B**). Since Ab blockade is not sufficient to eliminate all cytokine signaling, *Ifngr1^-/-^Il10^-/-^*mice were also generated and infected. Similar to Ab blockade, parasitemia is effectively controlled in *Ifngr1^-/-^Il10^-/-^*mice compared to *Il10^-/-^* controls, though it peaks later in the mice lacking IFN-γ signaling (**Fig. 6C,D**). Regardless, the overall parasite burden was significantly lower in *Ifngr1^-/-^Il10^-/-^*mice compared to WT mice. Additionally, blocking IFN-γ signaling in *Il10^-/-^* mice did not alter their ability to control parasite burden (**Fig. 6E,F**). Thus, these results suggest that the enhanced IFN-γ production associated with the loss of IL-10 signaling is not essential for improved control of parasite burden in *Il10^-/-^* mice.

**Fig 6.**
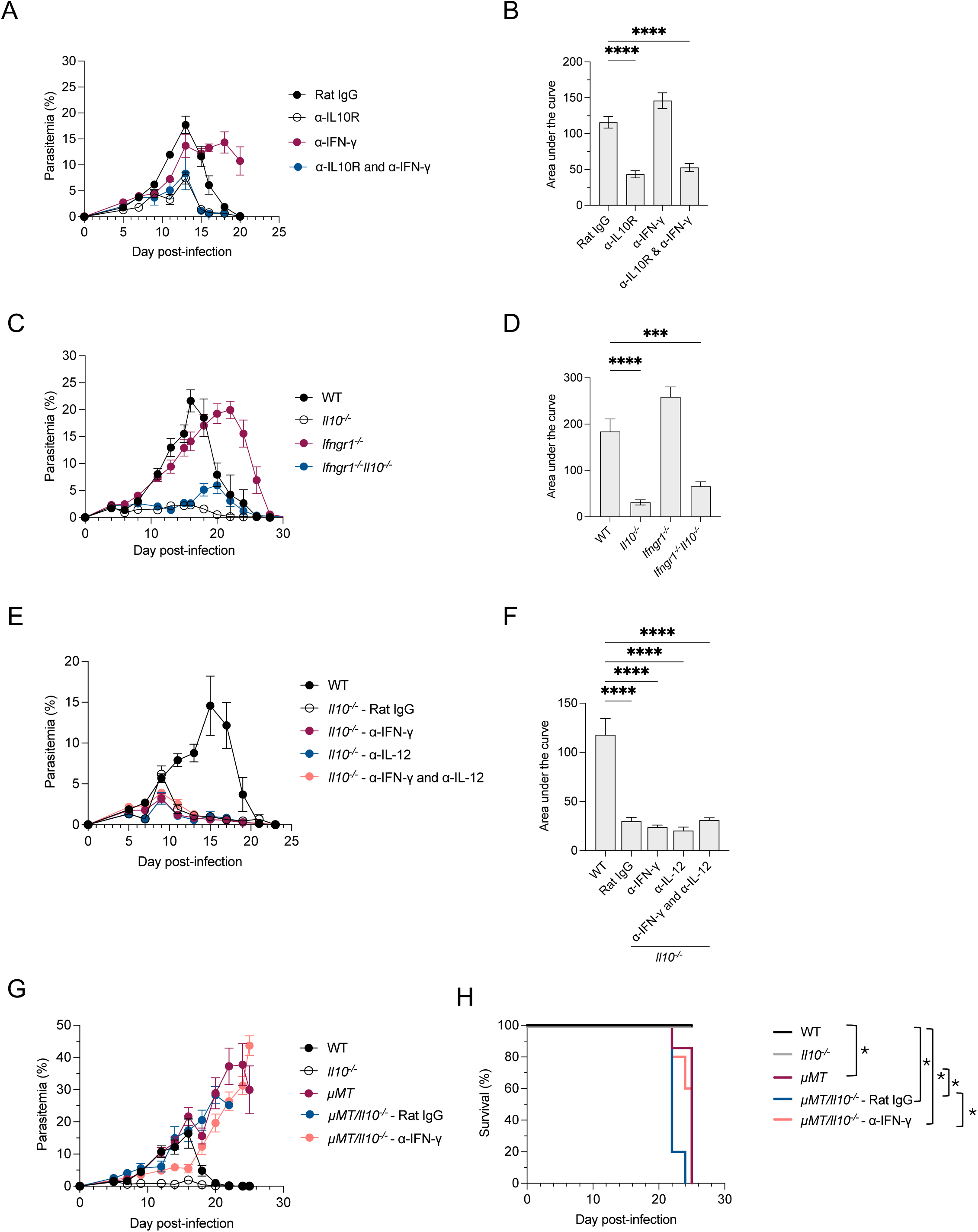
Enhanced IFN-γ or IL-12 does not contribute to protection in IL-10–deficient mice during *P. yoelii* infection. WT, *Il10^-/-^*, *μMT*, *μMTIl10^-/-^*, *Ifngr1^-/-^*, and *Ifngr1^-/-^Il10^-/-^* mice were infected i.p. with 10^5^ *P. yoelii* 17X pRBCs. (**A**) Representative parasitemia curve of the primary infection. WT mice were i.p. injected with either 200 μg of Rat IgG, 500 μg anti-IFN-γ, 200 μg anti-IL10R, or both anti-IFN-γ and anti-IL-10R at day 1 p.i. and every 3 days after. WT mice served as controls. (**B**) Area-under-the-curve analysis included data for the entire curve in A. (**C**) Representative parasitemia curve of the primary infection. (**D**) Area-under-the-curve analysis included data for the entire curve in C. (**E**) Representative parasitemia curve of the primary infection. *Il10^-/-^* mice were i.p. injected with either 500 μg of Rat IgG, 500 μg anti-IFN-γ, 500 μg anti-IL-12p75, or both anti-IFN-γ and anti-IL-12p75 at day 0, 3, 6, and 9 p.i. WT mice served as controls. (**F**) Area-under-the-curve analysis included data for the entire curve in E. (**G**) Representative parasitemia curve of the primary infection. *μMTIl10^-/-^*mice were i.p. injected with either 500 μg Rat IgG or 500 μg anti-IFN-γ at day 0, 3, 6, and 9 p.i. WT, *Il10^-/-^*, and *μMT* mice served as controls. **(H**) Representative survival curve for the primary infection in G. Mice were euthanized when the parasite burden exceeded 50% or when prominent signs of morbidity were observed. (**B, D, F**) A non-parametric Kruskal-Wallis test with a post hoc Dunn’s multiple comparisons test or **H**) a Mantel-Cox test determined significance. \**p* < 0.05, *** *p* < 0.001, **** *p* < 0.0001, ns = non-significant.

Since IL-12 production was also elevated in *Il10^-/-^*mice (**Fig. 2L, 3L**), and IL-12 has been implicated as a mediator of control in other *Plasmodium* models (45, 46), it was of interest to determine if this cytokine is critical for the improved protection observed in *Il10^-/-^* mice. Therefore, *Il10^-/-^* mice were given monoclonal Abs against IL-12 during *P. yoelii* infection. Similar to the results observed with anti-IFN-γ treatment in *Il10^-/-^* mice, administering anti-IL-12 to *Il10^-/-^* mice did not reduce their ability to control parasite burden compared with the Rat IgG control (**Fig. 6E-F**). To assess whether IFN-γ production compensates for the loss of IL-12 or vice versa, anti-IFN-γ and anti-IL-12 Abs were co-administered to *Il10^-/-^* mice during *P. yoelii* infection. As with individual treatments, co-administration of anti-IFN-γ and anti-IL-12 Abs in *Il10^-/-^* mice did not impair parasite control compared to the Rat IgG control (**Fig. 6E-F**). Therefore, these results suggest that neither IFN-γ nor IL-12 is necessary for *Il10^-/-^* mice to control *P. yoelii* infection.

Lastly, to determine if the B cell response compensates for the loss of IFN-γ, anti-IFN-γ monoclonal Abs were administered to *μMT*/*Il10^-/-^* mice during *P. yoelii* infection. IFN-γ blockade did not diminish parasite control or survival in the *μMT*/*Il10^-/-^*mice (**Fig. 6G, H**). In fact, parasite burdens and survival were slightly improved with IFN-γ blockade compared with the Rat IgG control (**Fig. 6G, H**). This suggests that, even in the absence of B cells, IFN-γ is not required for *Il10^-/-^*mice to maintain enhanced parasite control during the first two weeks of infection, compared with WT mice. Collectively, these results suggest that factors beyond B cells, IFN-γ, and IL-12 limit parasite expansion in *Il10^-/-^*mice for up to 14 days after infection with *P. yoelii.* After this point, B cells and their Ab product are necessary to maintain control of parasite burden and clear the infection.

## Discussion

Here, the results indicate that IL-10 production negatively impacts host-parasite control during *P. yoelii* infection, supporting some, but not all, previous findings (26). Whereas other studies have shown that mice treated with anti-IL-10R Abs exhibit increased susceptibility to *P. yoelii* infection and are unable to resolve it (42, 43), findings here indicate that IL-10-deficient mice and WT mice treated with anti-IL-10R Abs limit parasite burden more effectively than control mice without succumbing to the infection. The inability of C57BL/6 mice treated with anti-IL-10R Abs to control infection in the reported studies could stem from the dose and route of infection (10^6^ pRBCs i.v.), which differ from those used here. Alternatively, differences in the infectious tropism of the parasite strains could lead to greater invasion of normocytes than of reticulocytes. These differences might cause higher parasitemia and more severe anemia, and could trigger a stronger inflammatory response in the liver and/or lungs, resulting in severe immunopathology that could affect survival in the absence of IL-10. Although outcomes differ, the results here corroborate findings that IL-10 promotes humoral responses by aiding the accumulation of Tfh cells and GC B cells (42, 43). However, in their model, parasite-specific IgG responses decreased in the absence of IL-10; here, the absence of IL-10 did not affect parasite-specific IgG accumulation. It is possible that any defect in Ab production here was masked by higher serum IgG availability, given the lower parasite burdens in *Il10*^-/-^ mice infected with *P. yoelii*. Regardless, the Ab response generated in these *Il10*^-/-^ mice was sufficient to protect them against homologous rechallenge.

The finding that *Il10^-^*^/-^ mice survived infection with *P. yoelii* indicated that the immune response against the parasite was insufficient to induce severe tissue immunopathology, unlike in other *Plasmodium* infections. The absence of IL-10 is associated with hypoglycemia and hypothermia during the normally avirulent *P. chabaudi* infection (6, 37). It is also associated with cerebral complications, such as edema and hemorrhages, during *P. chabaudi*, *P. berghei*, and *P. falciparum* infections (14, 31, 47). Here, during *P. yoelii* infection, H&E staining confirmed that IL-10 prevents liver pathology in response to *P. yoelii* infection; the pathology observed in the absence of IL-10 was minimal and transient. More extensive hepatic necrosis and inflammation in the absence of IL-10 during *P. yoelii* infection has previously been reported (26). Nonetheless, findings here and by Couper et al. demonstrate that IL-10 is not critical for preventing lethal immunopathology (26).

IL-10 is implicated in preventing anemia (48), another disease pathology that can lead to complications or even death, during many human *Plasmodium* infections (31). Alternatively, IL-10 is also associated with promoting emergency myelopoiesis, leading to hematologic changes, including anemia and thrombocytopenia (16, 17, 49). Findings here demonstrate that anemia is more severe in the presence of IL-10 during *P. yoelii* infection, a finding that differs from previous reports indicating that IL-10 does not affect RBC counts during infection (26). Furthermore, given the role of IFN-γ in promoting hematopoietic changes (16, 50), it is surprising that the elevated systemic IFN-γ production observed in the absence of IL-10 does not lead to more severe anemia in *Il10*^-/-^ mice. Perhaps additional immunoregulatory mechanisms limit IFN-γ activity. The differences in RBC counts, hematocrit, and hemoglobin between WT and *Il10^-/-^* mice are most pronounced at peak parasitemia, suggesting that the exacerbated anemia in WT mice is due to elevated parasite burdens, which increase RBC lysis and turnover. Thus, the data here suggest that reducing parasite burden by suppressing IL-10 can reduce the severity of anemia during infection.

During the host response, IL-10 was shown to limit sustained T_H_1 polarization but did not suppress initial T_H_1 polarization. Although the T_H_1 response was prolonged in *Il10^-/-^* mice, with increased numbers of T-bet^+^ and CXCR6^+^ CD4^+^ T cells in the spleen through day 14 p.i., the enhanced IFN-γ production by CD4^+^ T cells was a transient phenotype associated with early infection. Furthermore, serum IFN-γ production steadily declined in *Il10^-/-^* mice, suggesting that factors other than IL-10 limit IFN-γ production by CD4^+^ T cells later in infection. These results are supported by previous work, showing that serum IFN-γ and TNF-α concentrations peak at day 7 p.i. but are then quickly downregulated in *Il10^-/-^*mice during *P. chabaudi* infection (37). Other potential anti-inflammatory mediators include TGF-β, which can limit circulating IFN-γ and TNF-α in *Il10^-/-^* mice during *P. chabaudi* infection (51). However, the mechanism by which IFN-γ is suppressed in *Il10^-/-^* mice during *P. yoelii* infection was not determined here and requires further investigation.

There was little evidence to suggest that an enhanced B cell response was mediating protection in the *Il10^-/-^* mice. Therefore, it was not surprising to find that B cells are not required during the first two weeks of infection for *Il10^-/-^* mice to maintain enhanced control of infection. This phenotype is consistent with previous work demonstrating that anti-IL-10R blockade in *μMT* mice reduces parasite burden during chronic *P. chabaudi* AS infection compared to control *μMT* mice (52). However, in the *P. chabaudi* model, blocking IL-10R signaling results in *μMT* mice succumbing to infection sooner than untreated *μMT* mice due to an enhanced pro-inflammatory response (52). Here, during *P. yoelii* infection, our findings corroborate an enhanced pro-inflammatory response in the *μMT/Il10^-/-^* mice. The relatively low parasite burdens at the time of death in the *μMT/Il10^-/-^* mice compared to the *μMT* mice suggest that the cause of death is likely an enhanced pro-inflammatory response leading to tissue pathology, or possibly exacerbated anemia. Previous reports indicate that the absence of B cells and IL-10 during *P. chabaudi* AS infection causes lethal tissue pathology (52), thus the extent of liver and lung damage in the *μMT/Il10^-/-^* mice during *P. yoelii* infection should be investigated further.

Although the mechanism of protection against acute parasitemia in *Il10*^-/-^ mice was B-cell-independent during the first half of the infection, the results here indicate that IFN-γ is also not required for protection in the absence of IL-10. Furthermore, even in the absence of a B cell response, the host is not reliant on IFN-γ to control parasitemia. These findings add to previous work in which simultaneous blockade of IL-10R and IFN-γ in C57BL/6 mice during *P. yoelii* infection significantly lowers parasite burden at day 14 p.i. (53). Moreover, the results here support data that suggest early protection against *P. yoelii* is independent of IFN-γ and the adaptive immune response (54). While our findings are supported by other studies in *P. yoelii*, they differ from previous reports utilizing the *P. chabaudi* model. During this infection, the loss of IL-10 production alone does not alter parasitemia, but elevated parasitemia was observed in *Ifngr1^-/-^Il10*^-/-^ mice (37), highlighting the importance of IFN-γ in mediating protection from acute parasitemia. Hence, infection of *Ifngr1^-/-^* mice results in elevated parasite burdens during *P. chabaudi* infection (37). In contrast, the results here indicate that *Ifngr1^-/-^Il10*^-/-^ mice can control their infection with *P. yoelii,* and the parasite burden resembles that observed in *Il10^-/-^*mice. Furthermore, mice deficient in IFN-γ signaling displayed a parasite burden similar to that of WT mice, though peak parasitemia was prolonged or delayed. These results suggest that IFN-γ may play different roles in controlling these infections, with its importance being more vital during *P. chabaudi* infection.

IL-12, a pro-inflammatory cytokine that promotes T_H_1 differentiation and subsequent IFN-γ production, was not vital to the protection afforded in *Il10^-/-^* mice. Blockade of IL-12 in WT mice has not previously been demonstrated during *P. yoelii* infection; however, previous reports indicate IL-12 is critical for protection from acute parasitemia during *P. chabaudi* and *P. berghei* infections (45, 46). In fact, the addition of recombinant IL-12 delays the onset of parasitemia during *P. berghei* infection (46). Thus, this provides further evidence for distinct protective mechanisms that control *P. yoelii* infection compared with other rodent-specific models.

Furthermore, the possibility of IL-12 or IFN-γ compensating for the loss of the other in IL-10–deficient mice was also ruled out. Changes in T_H_1 polarization in the absence of IL-12 and IFN-γ could be investigated in future studies to determine their impact on the T cell response. It’s possible that other cytokines, such as IL-27, still promote T-bet expression in CD4^+^ T cells in the absence of other T_H_1–polarizing cytokines (55). There could also be a shift toward more T_H_17 polarization. While there were very few IL-17^+^ CD4^+^ T cells in the spleen of WT or *Il10^-/-^* mice during *P. yoelii* infection (*data not shown*), the simultaneous loss of IL-12 and IFN-γ signaling could promote a shift in T cell polarization. Quantifying these populations, as well as subsequent neutrophil accumulation, is an important direction for future work.

While the results here have ruled out an early role for IL-12 and IFN-γ in parasite control in the absence of IL-10, this leaves one to ponder: if not these cytokines, then what is suppressing the early expansion of *P. yoelii* in these mice? Two possible candidates are IL-6 and TNF-α. Previous reports indicate that the injection of recombinant IL-6 reduces parasitemia and increases Ab production during *P. chabaudi* infection (56). Blocking IL-6 also results in elevated parasitemia late in *P. chabaudi* infection; however, it has little effect early in infection (57). Additionally, IL-6–deficient mice have elevated parasitemia during the first 14 days of infection with *P. yoelii* 17X (57). Thus, the trend of elevated IL-6 observed in the serum of *Il10^-/-^* mice early in infection could contribute to the more efficient parasite control seen in these mice. However, a previous report indicated that IL-6 promotes parasite control by enhancing the GC response and IgG class switching (57). If IL-6 plays a role in parasite control in *Il10^-/-^* mice, our data suggest it is not through this proposed mechanism, as GC B cell output is lower in *Il10^-/-^*mice than in WT controls. It was also reported that elevated serum TNF-α contributes to the susceptibility of IL-10–deficient mice to *P. chabaudi* infection, though blocking TNF-α in these mice reduced disease pathology without altering parasite burden (42). Given the results here of significantly elevated TNF-α in the *Il10^-/-^* mice early in infection, examining the importance of TNF-α in mediating parasite control in *Il10^-/-^* mice during *P. yoelii* infection through either Ab blockade or generation of *Tnfα^-/-^Il10^-/-^* mice is an important future direction.

Another possibility is that early IL-10 production alters the activity of innate cells in the spleen that recognize and clear iRBCs from circulation. The importance of phagocytic cells in controlling early parasite burden is illustrated by the finding that depletion of phagocytes with clodronate liposomes leads to an increase in parasite burden in response to infection with non-lethal and lethal strains of *P. yoelii* (54). Whether IL-10 modulates the expression of opsonic and non-opsonic receptors, such as CD36 (58), that are important for recognizing iRBCs in the spleen remains unknown. Alternatively, IL-10 could be affecting the production of free radicals or the expression of enzymes involved in parasite destruction. Another possibility is that IL-10 is promoting the development of myeloid suppressor cells, which can impair T and B cell responses to *Plasmodium* infection (59). If IL-10 is indeed impairing the early control of parasite burden by myeloid cells, understanding how it does so could offer a strategy to modulate its activity or enhance myeloid cell activity during this critical period of infection. Although blocking IL-10 activity during the first 2-4 days of a *P. yoelii* infection was shown to be detrimental to the outcome in one case (43).

In conclusion, IL-10 prevents control of *P. yoelii* infection independent of the B cell response, and of the T_H_1_-_associated cytokines IL-12 and IFN-γ. Future work aimed at further characterizing which cytokines and/or cell populations are critical for parasite control will inform the design of strategies to manipulate the host response and enhance protective immunity via vaccination or therapeutics, thereby reducing disease burden.

## Acknowledgements

*Plasmodium yoelii* 17X pRBCs were obtained through BEI Resources, NIAID, NIH: MRA-749, contributed by Dr. David Walliker. Recombinant MSP-1_19_ and AMA-1 proteins were obtained from Dr. James Burns Jr. (Drexel University College of Medicine). Anti-IFN-γ monoclonal Ab was provided by Dr. Christopher A. Hunter (University of Pennsylvania). We thank the Experimental Pathology Core at UAMS for their assistance with tissue sectioning and the H&E staining. We would also like to thank Andrea Harris from the Flow Cytometry core at UAMS for her assistance in sample processing.

## Author contribution

Conceived and designed the experiments: J.S.S. and M.D.J. Performed the experiments: M.D.J., K.A.O., A.R.G., S.L.Z., and J.S.S. Analyzed the data: M.D.J. and J.S.S. Wrote the paper: M.D.J. and J.S.S.

